# ActSeekN: A Structural-Motif–Based Pipeline for Interpretable Enzyme Function Annotation

**DOI:** 10.64898/2026.04.24.720574

**Authors:** Sandra Castillo, Chunhao Gu, Paula Jouhten, Gopal Peddinti, O. H. Samuli Ollila

**Affiliations:** Industrial Biotechnology and Food, VTT, Tekniikantie 21, 02044, Espoo, Otaniemi, Finland; Department of Bioproducts and Biosystems, Aalto University, Kemistintie 1, 02150, Espoo, Otaniemi, Finland; Institute of Biotechnology, University of Helsinki, Viikinkaari 5, 00014, Helsinki, Finland

**Keywords:** enzyme function annotation, structural motifs, active site identification, AlphaFold structures, EC number prediction

## Abstract

Accurate enzyme function annotation remains a major bottleneck in genome analysis despite the rapid expansion of available protein sequence and structure data. Most existing methods rely on sequence similarity or machine-learning representations, which often perform poorly for proteins with low sequence identity or convergent evolutionary histories. Because enzymatic activity is determined by the three-dimensional arrangement of catalytic and binding-site residues, structure-based approaches offer a mechanistically grounded alternative. However, their broader application has been constrained by the limited size and coverage of curated active-site reference databases. To address this challenge, we developed ActSeekN, a structural-motif–based functional annotation pipeline that combines the ActSeek active-site search algorithm with a newly constructed large-scale reference database derived from AlphaFold-predicted structures, UniProt annotations, and curated catalytic residue information. This framework enables rapid and scalable identification of conserved catalytic motifs across structurally related proteins, allowing function to be transferred on the basis of local three-dimensional catalytic geometry rather than global sequence similarity. In this way, ActSeekN overcomes a central limitation of previous structure-based methods by expanding the searchable space of catalytic motifs while retaining mechanistic interpretability. Benchmarking against state-of-the-art machine-learning approaches demonstrates competitive or superior performance. Applications to yeast, human, and Trichoderma reesei proteomes refine existing annotations, complete partial EC assignments, and identify previously unrecognized enzymatic functions, highlighting ActSeekN as a powerful tool for genome annotation and biotechnology.

**Graphical Abstract:** 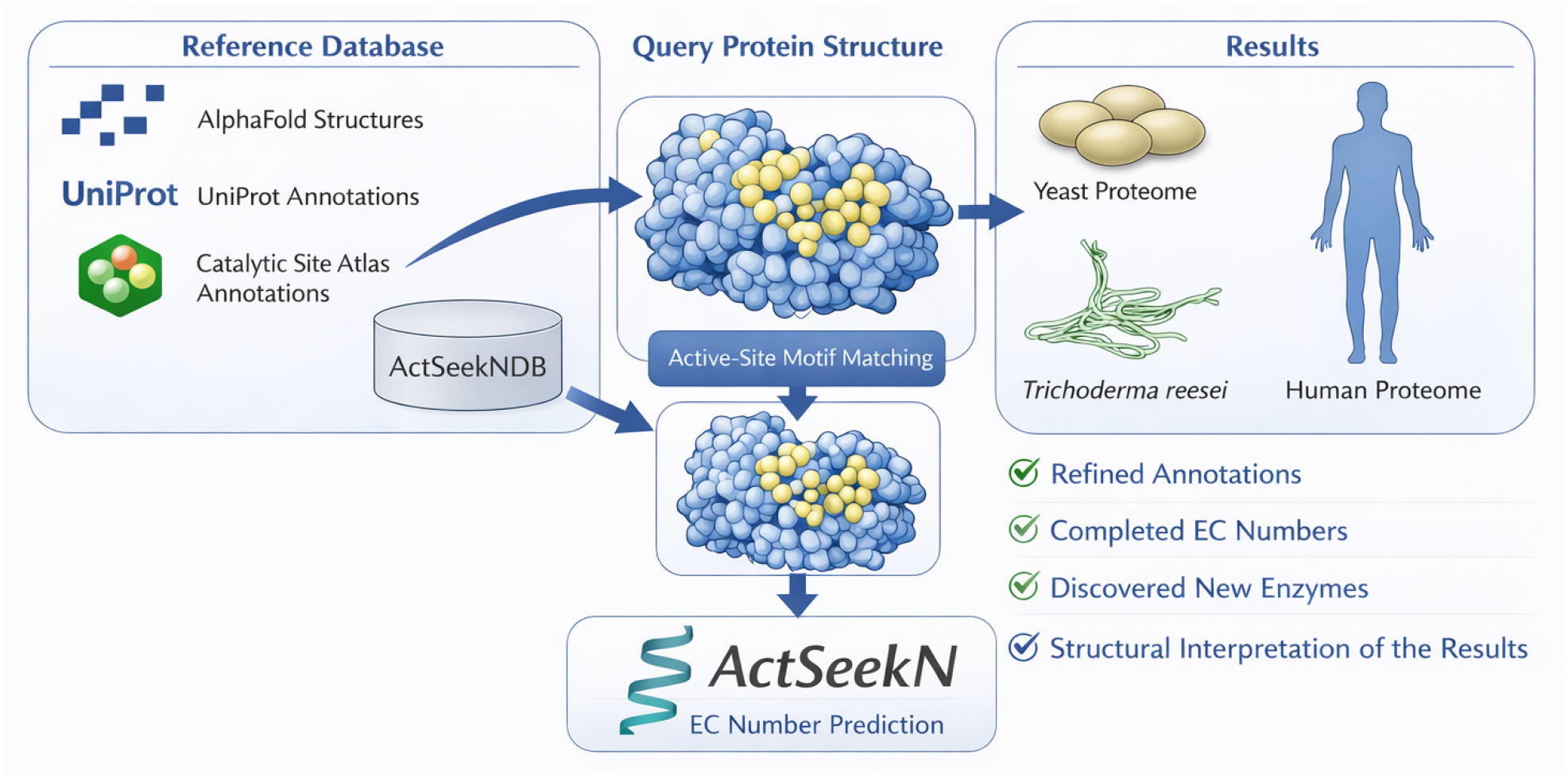

## Introduction

The remarkable diversity of enzyme functions is encoded in protein sequences, and advances in genome sequencing have made sequence data inexpensive to generate and abundant. However, the practical exploitation of this growing sequence space—both for understanding metabolism and for biotechnological applications—remains limited by the performance of computational methods for functional annotation. Accurately mapping protein sequences to specific biochemical activities is particularly challenging given the vast functional diversity of enzymes and the frequent absence of close homologs with experimentally characterized functions.

Most existing annotation methods rely primarily on sequence similarity to previously characterized proteins, under the assumption that homologous sequences perform similar functions. Orthology-based approaches, such as eggNOG-mapper (1), aim to improve function transfer by focusing on evolutionary conserved orthologs that are more likely to share function. In parallel, machine-learning approaches have been developed that learn functional patterns from large collections of amino-acid sequences or derived features, including deep learning methods (13; 15; 14). While these approaches have demonstrated impressive predictive performance, they typically operate as black-box models and provide limited mechanistic insight into why a particular function is assigned.

A key limitation shared by many sequence- and learning-based approaches is that enzyme function is not determined by global sequence similarity alone, but rather by the precise spatial arrangement of a small number of catalytic and binding-site residues. Enzymatic activity depends on the geometry, chemistry, and relative positioning of these residues, which together define the catalytic mechanism. As a result, proteins with low overall sequence identity—or those that have evolved convergently—may nevertheless share identical catalytic machinery, while closely related sequences can diverge functionally due to subtle changes in their active sites.

A small but important class of functional annotation methods explicitly exploits this principle by searching for conserved structural motifs corresponding to enzyme active or binding sites. These approaches not only assign functions but also identify the specific residues responsible for catalysis, offering mechanistic interpretability. Examples include GASS (10), COFACTOR (17) and EnzyMM (6). However, most structure-based motif search methods depend on curated databases of known active and binding sites, such as the Catalytic Site Atlas (CSA) (6). While highly reliable, these resources are inherently limited in scope: they contain annotations for only a few thousand enzymes.

These curated active-site databases are generated through labor-intensive processes that combine manual expert curation, experimental literature mining, and detailed structural analysis of individual enzymes. Consequently, their growth is slow and cannot keep pace with the rapid expansion of available sequence and structure data. This bottleneck has so far prevented structure-based annotation approaches from being used as general, proteome-scale functional annotation pipelines.

Recent breakthroughs in protein structure prediction, most notably AlphaFold (7), have dramatically changed the field by providing high-quality structural models for a large fraction of known proteins. Here, we used AlphaFold-predicted structures and curated UniProt (3) annotations to construct an extended database of enzyme structures with active sites and functions annotated. Integration of this database with the ActSeek enzyme mining algorithm (2) enables rapid, proteome-scale functional annotation by identifying conserved active-site geometries and transferring enzyme functions based on structural motifs rather than global sequence similarity. We demonstrated that this ActSeekN algorithm combined competitive predictive performance with mechanistic interpretability, allowing both accurate annotation and direct insight into the residues underlying enzymatic function. This overcomes the limitations imposed by small active-site databases and enables applying structure-based annotation at unprecedented scale and systematic detection of conserved catalytic geometries across diverse protein families and organisms.

## Materials and methods

### ActSeekNDB database development

The database includes 39761 proteins, each associated with one or several Enzyme Commission (EC) numbers. For each enzyme, we selected at least three amino acids that represent the active site of enzymes in that category.

We began by collecting all EC numbers listed in the UniProt database (3), totaling 6737 unique entries. For each EC number, we retrieved 250 proteins—both reviewed and unreviewed—from the UniProt database. These proteins were then grouped based on structural similarity using TM-align (18). To form clusters, we started with the first protein as the initial cluster representative. Each subsequent protein was aligned to this representative. If the root-mean-square deviation (RMSD) of the alignment was less than 0.3, the protein was added to the same cluster. Otherwise, a new cluster was created. When a protein could potentially belong to more than one cluster, it was assigned to the one with the lower RMSD. The number of proteins representing each EC number varied depending on the results of the structural clustering. Around 687 EC numbers could not be included due to the absence of structural data in the AlphaFold database or annotations.

For clusters containing more than two proteins, conserved amino acids were identified through structural alignment. One representative protein from each cluster was then selected randomly for inclusion in the database. Active-site and binding-site annotations were assigned by intersecting UniProt-annotated residues across all proteins within a cluster and combining them with the structurally conserved residues. In cases where no UniProt annotations were available, an AI-based agent (based on ChatGPT5.1 (11)) was used to assist in annotation. The agent selected three amino acids located between conserved residues that appeared spatially clustered by integrating multiple sources of prior information, including the atomic coordinates, identities and positions of conserved residues, the EC number associated with the protein, and curated annotations of active-site and binding-site residues from the UniProt database. This contextual information guided the agent to select residues probably located within functionally relevant regions of the catalytic site. The number of proteins representing each EC number varied depending on the outcome of the structural clustering.

In addition, we integrated data from the Catalytic Site Atlas, retaining the original annotations and using three annotated amino acids as the primary search motifs. However, this resource often provided fewer than the required three annotated residues for the active site. In such cases, missing residues were supplemented using conserved amino acids identified through structural alignment.

Each protein in the ActSeekNDB was assigned a confidence score ranging from 0 to 5, derived from the UniProt annotation score associated with that protein. A score of 0 was assigned to proteins that have been removed from the UniProt database at the last database update. Importantly, this confidence score does not measure the correctness of the functional annotation predicted by ActSeekN. Instead, it reflects the reliability of the reference protein used as the source for annotation transfer. Higher scores correspond to proteins with stronger experimental or curated evidence in UniProt, whereas lower scores indicate proteins whose annotations are based on limited evidence, computational inference, or homology. Consequently, the confidence score provides an estimate of how trustworthy the annotation source is, rather than an assessment of the predicted function itself.

### ActSeekN algorithm

The standalone ActSeekN tool, which combines the previously published ActSeek structural motif search algorithm (2) with the ActSeekNDB reference database, was implemented in standard C++. The protein coordinates and distance calculations were handled using the C++ linear algebra library Eigen. Multi-threaded comparisons against seed proteins were supported via the `<thread>` library. Efficient data sharing between processes was implemented using the Boost Interprocess library (`<boost/interprocess>`). Construction of KD-trees and nearest-neighbor searches were performed using the nanoflann C++ library. ActSeekN additionally provides optional GPU acceleration via CUDA, enabling distance calculations to be offloaded to NVIDIA GPUs when available, thereby improving performance for large-scale analyses. The software is distributed as an open-source standalone tool and is publicly available at https://github.com/Aalto-Microbial-Physiology/ActSeekN. Across the Price dataset (15), the average running time per protein is 15 seconds, with execution times ranging from few seconds to one minute depending primarily on the size of the query protein.

### Web Application

To enable easy use of ActSeekN, a web interface is provided at https://actseek.vtt.fi. The web application was developed using Django and hosted using nginx on a virtual machine equipped with 8 CPUs and 32 GB RAM, running `Ubuntu 24.04 LTS`. It allows the users to query using PDB files or UniProt Accession Numbers, limiting the maximum number of input entries to 100 in a single query. When a query contains multiple inputs, the server processes them asynchronously and provides the results on a single page that gets updated progressively as individual results become available. The results page URL can be bookmarked, the browser window closed and revisited later for viewing the results.

### Evaluation of annotations

We evaluated our algorithm using the same datasets as the machine-learning–based CLEAN-contact method: the New392 dataset, which contains 392 proteins, and the Price dataset, which contains 195 proteins. When the protein providing the functional annotation was identical to the query protein present in the dataset, that protein was excluded from the results, and only the remaining hits were considered.

To ensure that the comparison with previous annotation methods was done fairly, we used the same evaluation metrics and code (available at (16)) as in CLEAN-Contact (14). For ActSeekN, only the top five EC numbers were retained for annotation, and EC numbers associated with less than 0.15% residue position similarity were filtered out. Performance was assessed using precision, recall, F1-score, and the area under the curve (AUC). Precision measures the proportion of predicted positive annotations that are correct, reflecting the reliability of the predictions. Recall quantifies the proportion of true positive annotations that are successfully identified, capturing the method’s ability to recover relevant functional relationships. The F1-score, defined as the harmonic mean of precision and recall, provides a single summary measure that balances these two complementary aspects of performance. Finally, AUC evaluates the model’s ability to discriminate between positive and negative classes across all possible decision thresholds, with higher values indicating better overall ranking performance independent of a specific cutoff.

## Results

### ActSeekN annotation strategy

ActSeekN annotation strategy, illustrated in Fig. 1, is based on previously published ActSeek program (2) and ActSeekN database (ActSeekNDB) developed here. ActSeekNDB contains representative structures and active site amino acids for EC numbers related to known enzyme functions (Fig. 1 A). Active sites in the protein to be annotated (query protein) are then identified using ActSeek search with each active site defined in ActSeekNDB as a seed and the query protein as the target (Fig. 1 B). As a result, the algorithm identifies all active sites present in the query protein (Fig. 1 C). Because these active sites are associated with protein functions in ActSeekNDB, this enables the functional annotation of query protein functions.

**Figure 1.**
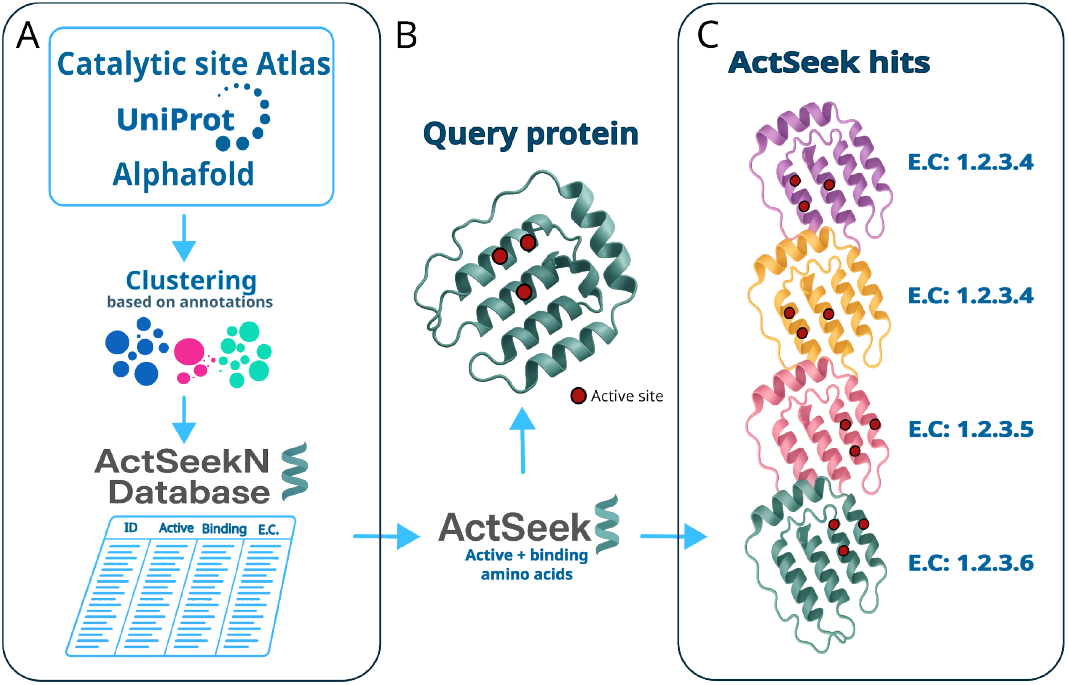
Schematic overview of the enzyme annotation workflow. **A.** ActSeekN database is built by clustering annotated UniProt structures and integrating the Catalytic Site Atlas. **B**. Structural motifs in the query protein are identified using ActSeek algorithm with the ActSeekN database. **C** Identified functions are annotated to the query protein .

ActSeekNDB currently contains 39761 proteins, each associated with active site residues and functionalities dispersed into 6050 EC numbers. The database was composed by first clustering proteins structures from UniProt (3), selecting representative structures from each cluster, adding a consensus annotation of the active or binding site amino acids and finally adding the proteins from the Catalytic Site Atlas database with known EC numbers. The current distribution of functions in the ActSeekNDB database is shown in figure 2. ActSeekNDB is regularly updated following the changes in UniProt. Therefore, the number of structures and EC number distributions in ActSeekNDB are evolving with time.

**Figure 2.**
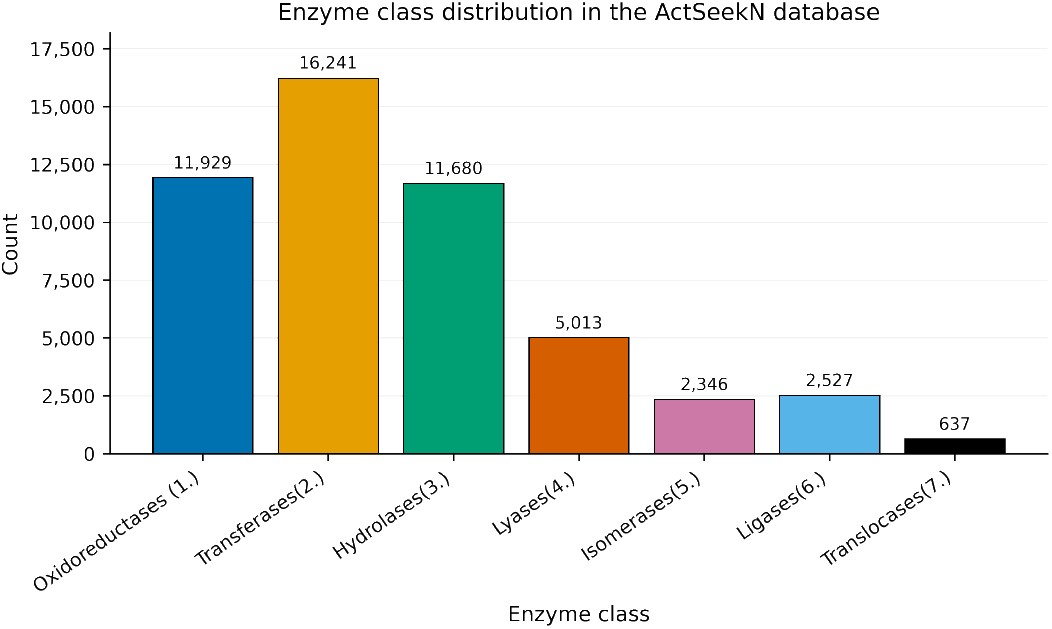
Barplot of the EC number distribution in the ActSeekNDB database.

For the functional annotation of a given organism, the search described in Fig. 1 was run with all relevant proteins from the organism as queries. This resulted a list of functions in terms of EC numbers with different confidence levels for each annotated protein.

### Benchmark results

We benchmarked ActSeekN using the New-392 and Price-149 datasets that were originally introduced to test the performance of CLEAN (15) and later used to benchmark also the CLEAN-Contact (14). The New-392 dataset contains 392 enzyme sequences distributed over 177 different EC numbers and Price-149 comprises 149 enzyme sequences distributed over 56 different EC numbers. Because CLEAN-Contact outperformed other annotation tools in previous tests (14; 13), we benchmarked ActSeekN against CLEAN-Contact using the same procedure as was used in their publication (14).

Benchmarking results in Fig. 3 suggested that ActSeekN outperformed CLEAN-Contact in Precision, Recall and F1-score when evaluated against New-392 dataset and Price-149. Additionally, ActSeekN provides users with the rationale behind each selected annotation, as well as information on potentially active or binding sites.

**Figure 3.**
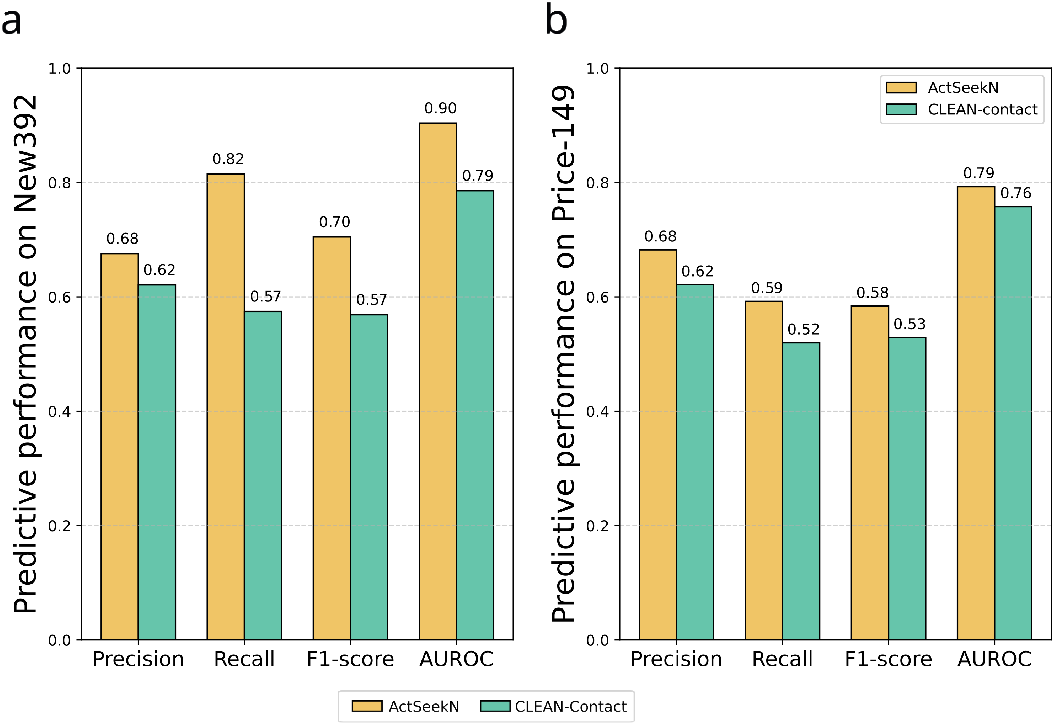
Comparison of predictive performance between ActSeekN and CLEAN-Contact measured by Precision, Recall and F1-score on the New-392 and Price-149 dataset.

### Functional annotation of yeast

To demonstrate that ActSeekN can to improve functional annotation of real organisms, we applied it to a common eukaryotic model and application workhorse, yeast *Saccharomyces cerevisiae* (reference strain S288C). ActSeekN achieved an overall prediction accuracy of 88.9%. ActSeekN completed 192 previously incomplete EC numbers. For genes lacking EC annotations in UniProt but associated with Gene Ontology (GO) molecular function terms (324 genes), the predicted EC numbers were evaluated for consistency by comparing the enzymatic activities implied by the EC assignments with the corresponding GO functional descriptions. In most cases, the assigned EC numbers matched the known or expected molecular functions of the proteins. Representative cases of these updates—including molecular chaperones, transporters, metabolic enzymes, and previously uncharacterized proteins—are summarized in Table 1. This table highlights how ActSeekN’s predictions align with or refine existing functional annotations, offering new enzymatic insights for several proteins lacking experimental characterization.

**Table 1.**
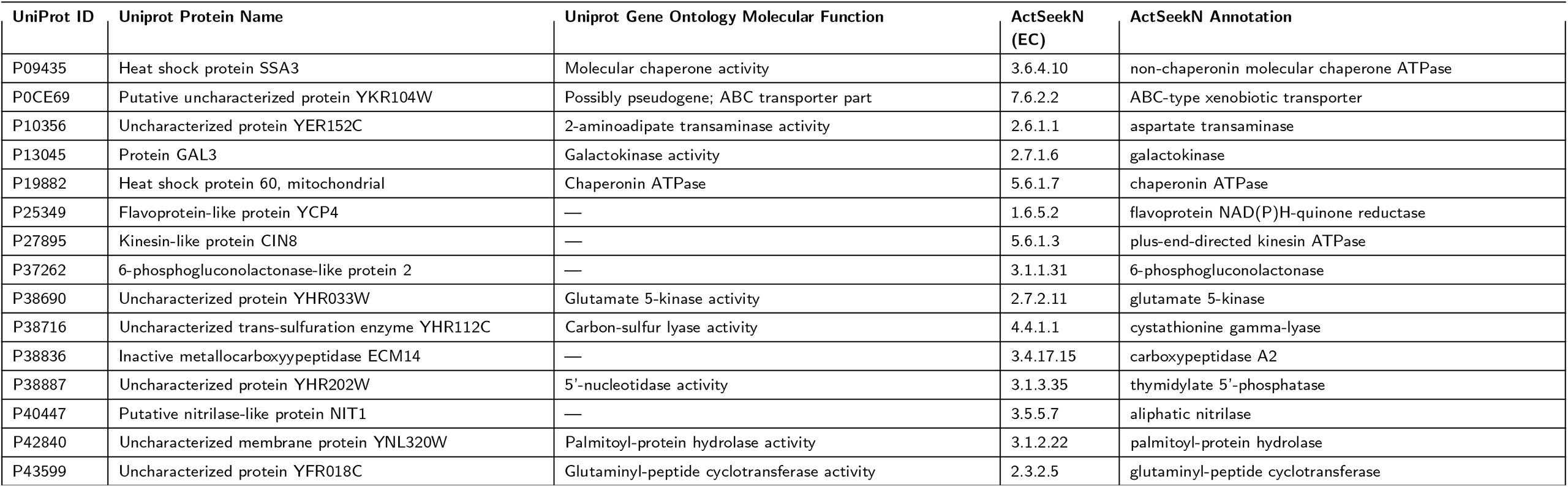
Examples of enzyme annotations generated by ActSeekN for *S. cerevisiae* proteins lacking EC numbers in UniProt. For each enzyme, the predicted EC number is shown together with the corresponding Gene Ontology (GO) molecular function or the protein name, illustrating consistency between predicted enzymatic activity and known functional descriptions.

In some cases, the function assigned by ActSeekN did not match the UniProt annotation, or the true function of the gene remains unknown. A notable example is the protein P40037 associated with the HMF1 gene (YER057C). It is originally annotated as a high-dosage growth inhibitor, whereas ActSeekN predicted it to be a 2-iminobutanoate/2-iminopropanoate deaminase (EC 3.5.99.10). Structural superposition of P40037 protein with the representative enzyme for EC 3.5.99.10 in the ActSeekNDB revealed that both share identical catalytic amino acids and a perfectly matched overall fold (Fig. 4). Similarly, the protein P42840 associated with gene N0342 (YNL320W), currently annotated in UniProt as an uncharacterized membrane protein, was assigned by ActSeekN as a palmitoyl-protein thioesterase (ABHD10, EC 3.1.2.22). Although structural superposition showed that P42840 did not align perfectly with the representative structure for EC 3.1.2.22 in the ActSeekNDB, the key active and binding-site residues matched in both position and amino acid identity (Figure 4).

**Figure 4.**
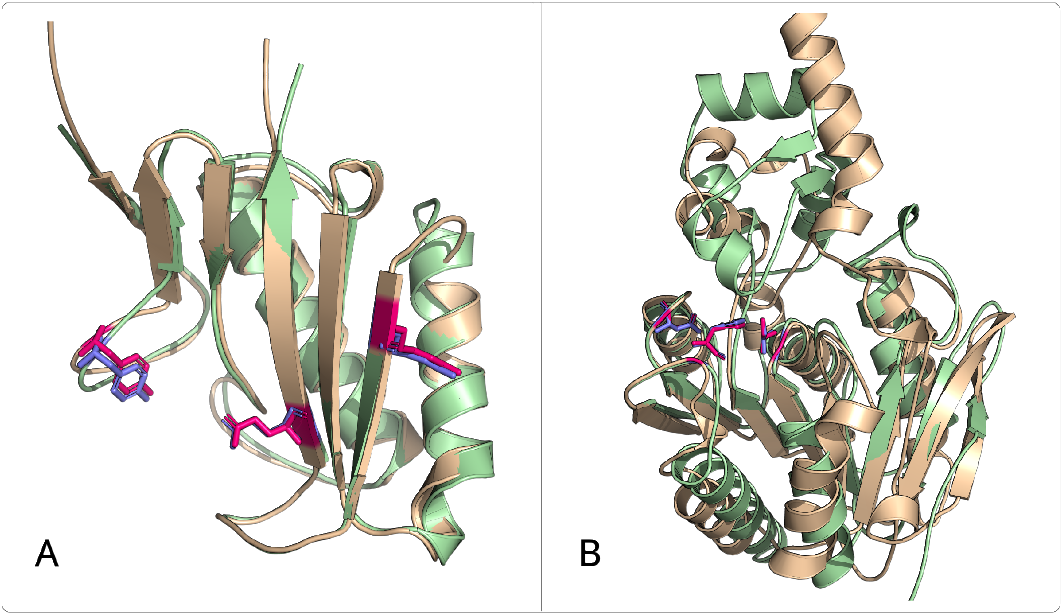
**A**. Structural superposition (PyMOL) of the *S. cerevisiae* protein P40037 (HMF1 / YER057C), annotated as a high-dosage growth inhibitor, with the representative protein for EC 3.5.99.10 in the ActSeekNDB, a 2-iminobutanoate/2-iminopropanoate deaminase from *E. coli*. Active and binding-site residues are highlighted in red for the yeast protein (colored wheat) and blue for the *E. coli* protein (colored light green). **B**. Structural superposition of the *S. cerevisiae* protein P42840 (N0342 / YNL320W), annotated as an uncharacterized membrane protein, with the representative protein for EC 3.1.2.22 in the ActSeekNDB, a palmitoyl-protein thioesterase protein Q5E9H9 from *B. taurus*. The three active-site residues are marked in red for the yeast protein (colored wheat) and blue for the *B. taurus* protein (colored light green).

These findings highlighted the value of computational annotation tools like ActSeekN in refining and correcting existing protein function annotations, even in well-studied organisms such as *S. cerevisiae*.

### Functional annotation of human

As a further test of the ActSeekN, we applied it to the human genome. Overall, 90% of ActSeekN predictions matched the EC numbers already assigned in UniProt, indicating high accuracy. Among these, 521 enzymes had incomplete EC annotations in UniProt, for which ActSeekN inferred more specific enzymatic functions. Despite the lack of formal EC number annotations, many of these enzymes have a functional descriptions in Uniprot. ActSeekN assigned EC numbers were mostly consistent with these descriptions as exemplified in table 2.

**Table 2.**
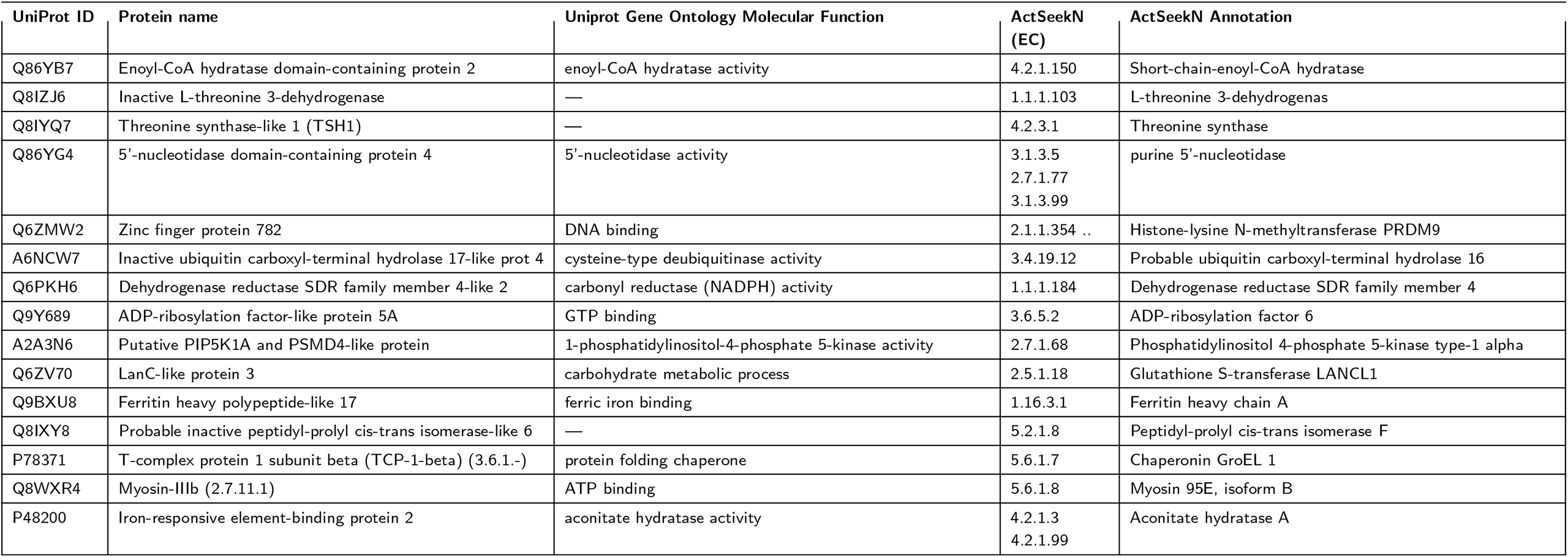
Examples of enzyme annotations generated by ActSeekN for human proteins lacking EC numbers in UniProt. For each enzyme, the predicted EC number is shown together with the corresponding Gene Ontology (GO) molecular function or the protein name, illustrating consistency between predicted enzymatic activity and known functional descriptions.

In cases where the existing functional description disagreed with the ActSeekN annotation, the tool usually reported a low confidence score indicating that ActSeekN could correctly evaluate its potentially spurious annotations. Nevertheless, we further investigated two examples where ActSeekN provided a high-confidence annotation despite these discrepancies. While *SLC3A1* (Uniprot: Q07837) is conventionally annotated as an amino acid transporter heavy chain, the ActSeekN pipeline identified it as a trehalose-6-phosphate hydrolase (EC 3.2.1.93). Structural superposition of Q07837 with the representative enzyme for EC 3.2.1.93 in the ActSeekNDB (*E. coli treC* ; Uniprot: P28904) yielded a Root Mean Square Deviation (RMSD) of 1.1 Å, revealing high structural conservation and the retention of three key catalytic residues (Figure 5A). However, hydrolase activity has not been reported for *SLC3A1* (Uniprot: Q07837). This discrepancy likely reflects an evolutionary functional divergence; although the ancestral glycosyl hydrolase scaffold and primary active site residues remain intact, subtle structural substitutions may have abolished hydrolase activity in favor of its contemporary role in mammalian amino acid transport.

**Figure 5.**
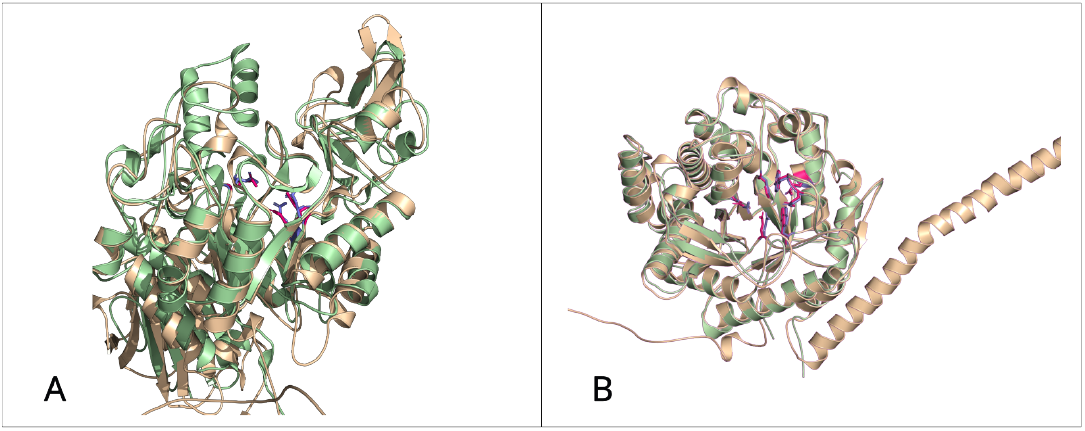
**A**. Superposition of the human *SLC3A1* protein (Uniprot: Q07837) commonly annotated as amino acid transporter (wheat color) with the *E. coli treC* protein (Uniprot: P28904), which is the representative enzyme for trehalose-6-phosphate hydrolases (EC 3.2.1.93); *E. coli treC* ; Uniprot: P28904) in the ActSeekNDB (light green). Catalytic amino acids are marked in red for the human protein and in blue for the *E*.*coli* protein.**B**. Superposition of the human *LCTL* protein commonly annotated as lactase-phlorizin hydrolase (wheat color) with the representative enzyme for *β*-glucosidase (EC 3.2.1.21); human GBA3; UniProt: Q9H227) in the ActSeekNDB (light green). Catalytic amino acids are marked in red for *LCTLn* and in blue for *GBA3*.

The *LCTL* gene (UniProt: Q6UWM7) is annotated as a lactase-like protein and has been suggested to play a structural or enzymatic role in the ocular lens suture, although this function has not been experimentally validated. LCTL exhibits high sequence homology to domain IV of lactase-phlorizin hydrolase (*LCT*), while lacking the other domains present in the full-length lactase protein. Functional annotation via ActSeekN identified *LCTL* as a putative *β*-glucosidase (EC 3.2.1.21), showing high homology to the representative structure in ActSeekNDB for that EC number (human cytosolic *β*-glucosidase; GBA3; UniProt: Q9H227) (Figure 5B). Alignment of their catalytic domains revealed that all critical binding and active-site residues are conserved, with the exception of a single conservative substitution where Glutamate is replaced by Aspartate. Given that both residues contain carboxylate side chains capable of serving as general acid/base catalysts, this substitution may allow retention of *β*-glucosidase activity, while potentially affecting substrate specificity through modifications to the catalytic region (5; 4).

### *Trichoderma reesei* proteases

*Trichoderma reesei* is a well-established filamentous fungus widely used in industrial biotechnology, particularly for the production of heterologous proteins. Its natural ability to secrete large amounts of enzymes, especially cellulases (9), makes it an attractive host for protein expression. Over the years, it has been genetically engineered to enhance its secretion capacity, reduce unwanted metabolic pathways, and improve overall yield (12).

One of the major challenges in using *T. reesei* for heterologous protein production is the degradation of target proteins by endogenous proteases. These proteases can act in the production process, significantly reducing the yield and stability of the desired proteins. To address this, many protease genes have been systematically deleted or silenced in production strains to minimize proteolytic activity and improve product stability (8).

Approximately 75% of the *T. reesei* genes were annotated according to the Uniprot annotations, being less extensively studied than *S. cerevisiae* and human. Furthermore, many UniProt annotations rely primarily on sequence similarity rather than experimental evidence. Nevertheless, ActSeekN provided EC number annotations for more than 2,000 proteins that were unannotated in UniProt.

Given the impact of proteases on production efficiency, we focused here on newly annotated *T. reesei* proteases. ActSeekN annotated 182 protease-encoding proteins in *T. reesei* with EC number “3.4.-.-”, which was significantly more than the 64 proteases currently annotated in the UniProt database. Similarly to yeast and human, most proteases identified by ActSeekN that lack EC number annotations in UniProt possessed Gene Ontology (GO) annotations consistent with protease activity suggested by ActSeekN. Nevertheless, in some cases ActSeekN predicted protease function whereas the corresponding GO annotations were absent or inconsistent. For example, the *T. reesei* protein with UniProt ID G0RSP4 lacked UniProt annotation but was classified by ActSeekN as a proline iminopeptidase with the EC number of 3.4.11.5 (Figure 6A). Similarly, the protein with UniProt ID G0RAG5 had no GO annotation in UniProt, yet ActSeekN annotated it as a Gamma-glutamyl peptidase 1 with EC number 3.4.19.16 (Figure 6B).

**Figure 6.**
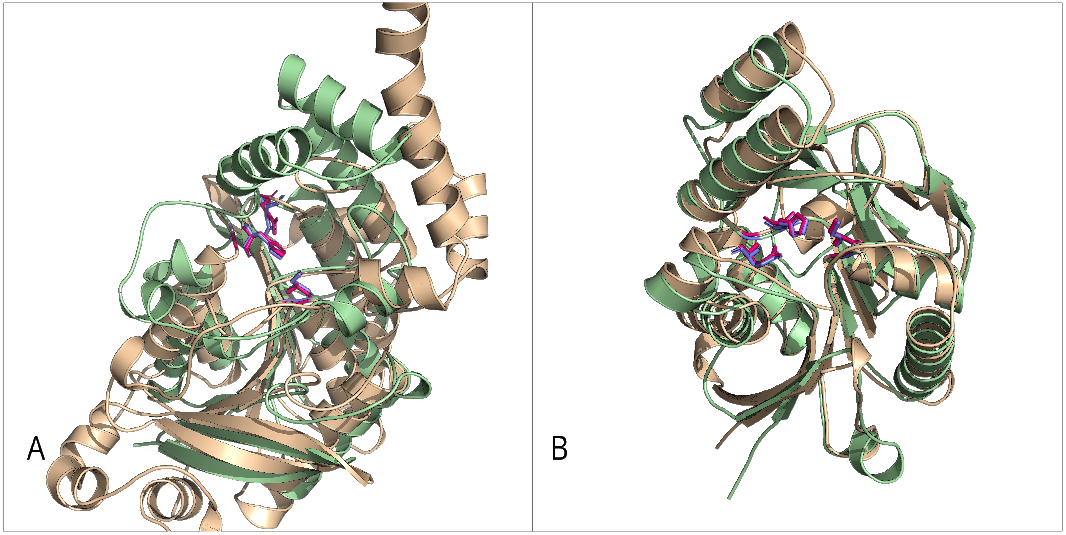
**A**. Superposition of the *T*.*reesei* protein G0RSP4 (wheat color) with the representative protease for EC number 3.4.11.5 (*Heyndrickxia coagulans* protein *pip*; Uniprot: P46541) in the ActSeekNDB (light green). Catalytic amino acids are marked in red for the *T*.*reesei* protein and in blue for the *H. coagulans* protein. **B**. Superposition of the *T*.*reesei* protein G0RAG5 (wheat color) with the representative protease for EC number 3.4.19.16 (*Arabidopsis thaliana* protein *GGP1* ; Uniprot:Q9M0A7) in the ActSeekNDB (light green). Catalytic amino acids are marked in red for the *T*.*reesei* protein and in blue for the *A. thaliana* protein

This expanded identification provides a valuable resource for further strain engineering and optimization of *T. reesei* as a robust platform for industrial protein production.

## Discussion

In this work, we presented ActSeekN, a structural-motif–based functional annotation pipeline that leverages curated active- and binding-site geometries to assign enzyme functions. By focusing on the spatial arrangement of catalytic residues rather than overall sequence similarity or black-box feature embeddings, ActSeekN addresses a key limitation of many existing annotation approaches: their difficulty in assigning the same biochemical function to proteins that share low sequence identity or have evolved similar activities through different evolutionary paths.

A major strength of ActSeekN lies in the combination of the previously published ActSeek enzyme mining tool (2) and ActSeekNDB developed here. ActSeekNDB integrates annotations from UniProt and the Catalytic Site Atlas with structurally conserved residues identified through clustering of AlphaFold models and UniProt annotations. The ActSeek mining algorithm is then used together with ActSeekNDB to identify candidate active-site residues associated with different enzyme functions in a query protein. This strategy enables ActSeekN to capture mechanistically relevant information even when sequence similarity is low and to provide interpretable annotations grounded in specific catalytic or binding-site residues. Recent structure-based learning approaches such as TopEC (13) have shown that incorporating localized three-dimensional descriptors can improve robustness to fold and sequence diversity, although their overall EC classification performance remains slightly below that of sequence-based methods such as CLEAN (15). Importantly, these machine-learning approaches infer function from learned representations and do not explicitly identify catalytic residues or active sites. In contrast, ActSeekN directly links functional assignments to specific residue-level annotations, offering insight into *why* a particular function is assigned and facilitating manual validation and hypothesis generation. CLEAN-Contact (14) similarly improves prediction accuracy by incorporating structural context, but does not yield explicit mechanistic annotations.

Benchmarking against CLEAN-Contact on the New-392 and Price datasets demonstrated that ActSeekN achieved competitive—and in these datasets superior—performance in terms of precision, recall, and F1-score. Importantly, these gains are not achieved at the expense of interpretability. ActSeekN additionally reports the specific residues and local structural context underlying each annotation, which can be particularly valuable when evaluating borderline predictions or exploring alternative functional hypotheses. This transparency is especially relevant in cases where multiple EC numbers are plausible or where UniProt annotations are incomplete or inconsistent.

Application of ActSeekN to the yeast and human proteomes highlights its practical utility in refining existing annotations and proposing new enzymatic functions for proteins lacking complete EC numbers. In well-studied organisms such as *S. cerevisiae*, ActSeekN was able to complete or refine many incomplete EC annotations and to assign functions consistent with Gene Ontology molecular function terms. In humans, ActSeekN frequently assigned plausible enzymatic activities to proteins annotated with broad or non-enzymatic functional descriptions, while appropriately reporting low confidence scores when predicted activities conflicted with existing annotations. Detailed case studies, including HMF1 in yeast and LCTL in humans, illustrate that conserved catalytic geometries can enable accurate identification of enzymatic activity even when sequence similarity or annotated biological roles differ, highlighting the power of structure-based annotation. Since conserved catalytic motifs do not necessarily indicate conservation of the protein’s full biological function or physiological role, these findings may require further study for gaining a more accurate understanding of the functions of these proteins.

ActSeekN is not intended to replace experimental validation or comprehensive functional characterization. The presence of a conserved catalytic motif does not guarantee enzymatic activity *in vivo*, particularly in cases of regulatory inactivation, altered substrate specificity, or repurposing of enzymatic scaffolds for non-catalytic roles. This is exemplified by proteins that retain active-site residues but may no longer perform the corresponding reaction due to subtle structural changes outside the core motif, such as human *SLC3A1* (Uniprot: Q07837). For this reason, ActSeekN confidence scores and supporting structural evidence should be interpreted as indicators of functional plausibility rather than definitive proof of activity.

Several limitations of the current implementation should be acknowledged. First, ActSeekNDB coverage depends on the availability and quality of structural models and curated annotations. Although AlphaFold has dramatically expanded structural coverage, certain EC numbers could not be included due to missing or low-confidence models. Second, the requirement for at least three spatially conserved residues may limit sensitivity for enzyme classes with poorly characterized or highly flexible active sites. Third, while ActSeekN scales efficiently to proteome-wide analyses, runtime remains dependent on protein size and motif complexity, going from less than 2 second to few minutes.

Future developments will focus on expanding ActSeekNDB coverage as UniProt and structural databases continue to evolve, refining confidence estimation by integrating additional structural and evolutionary features, and combining ActSeekN with complementary machine-learning approaches. Hybrid pipelines, potentially powered by AI-agents, that use ActSeekN for high-confidence, interpretable annotations and ML-based methods for broader functional prioritization may offer a particularly powerful strategy. In addition, systematic experimental validation of selected high-confidence predictions will be essential to further assess the strengths and limitations of structural motif–based functional annotation.

Overall, ActSeekN provides a robust, interpretable, and scalable framework for enzyme function annotation that complements existing sequence- and machine-learning–based approaches. By explicitly linking predicted functions to conserved catalytic geometry, it offers both practical annotation capabilities and mechanistic insight, making it a valuable tool for genome annotation, enzyme discovery, metabolic modeling and hypothesis-driven experimental design.

## Conclusions

In this study, we presented ActSeekN, a structure-based functional annotation framework that infers enzyme function at proteome scale by exploiting conserved catalytic and binding-site geometries. By focusing on the spatial arrangement of functionally important residues rather than global sequence similarity, ActSeekN complements sequence-based and machine-learning methods while providing mechanistically interpretable predictions.

Benchmarking against the state-of-the-art CLEAN-Contact method showed that ActSeekN achieves improved predictive performance on widely used datasets and provides explicit structural evidence for each annotation. This interpretability helps users to distinguish high-confidence functional assignments from weaker, hypothesis-generating predictions, and supports more targeted protein engineering.

Applications to the yeast, human, and *T. reesei* proteomes demonstrated ActSeekN’s utility for refining annotations, completing incomplete EC numbers, and identifying functions for previously uncharacterized proteins. ActSeekN also reveals that conserved catalytic motifs can persist despite evolutionary functional divergence, underscoring both the value and the limitations of structure-based annotation transfer. In *T. reesei*, the expanded identification of proteases provided useful insights for strain engineering and process optimization.

Although ActSeekN does not replace experimental validation, it offers a robust and transparent framework for prioritizing functional hypotheses and guiding downstream analyses. As structural databases and curated annotations continue to grow, ActSeekNDB can be updated to improve coverage and accuracy. Overall, ActSeekN provides a scalable, interpretable, and biologically grounded approach to enzyme function annotation with broad applications in genome annotation, enzyme discovery, metabolic modeling, and biotechnology.

## Abbreviations

EC: GO
CSA: RMSD

## Acknowledgements

We thank to Alejandro Revuelta for his valuable contribution on the conceptualization of the standalone application.

Author contributions: Conceptualization SC and OHSO; Database Development SC; Standalone Application Development SC, CG and PJ; Web Application Development GP; Writing Original Draft SC, OHSO, CG, PJ and GP.

## Confict of interest

All authors declare that they have no conflicts of interest.

## Data availability

ActSeekN is available as a standalone package via the GitHub repository https://github.com/Aalto-Microbial-Physiology/ActSeekN and can also be accessed through a web interface at https://actseek.vtt.fi.

## References

1. Carlos P Cantalapiedra, Ana Hernández-Plaza, Ivica Letunic, Peer Bork, and Jaime Huerta-Cepas. eggnog-mapper v2: functional annotation, orthology assignments, and domain prediction at the metagenomic scale. Molecular biology and evolution, 38(12):5825–5829, 2021.

2. Sandra Castillo and Osmo Henri Samuli Ollila. Actseek: fast and accurate search algorithm of active sites in alphafold database. Bioinformatics, 41(8):btaf424, 07 2025.

3. The UniProt Consortium. Uniprot: the universal protein knowledgebase in Nucleic acids research, 2025. 53(D1):D609–D617, 2025.

4. Erik W. Debler, Kanishk Jain, Rebeccah A. Warmack, You Feng, Steven G. Clarke, Günter Blobel, and Pete Stavropoulos. A glutamate/aspartate switch controls product specificity in a protein arginine methyltransferase. Proceedings of the National Academy of Sciences, 113(8):2068–2073, 2016.

5. Tracey M. Gloster, Johan P. Turkenburg, Jennifer R. Potts, Bernard Henrissat, and Gideon J. Davies. Divergence of catalytic mechanism within a glycosidase family provides insight into evolution of carbohydrate metabolism by human gut flora. Chemistry & Biology, 15(10):1058–1067, 2008.

6. Raymund E. Hackett, Ioannis G. Riziotis, Martin Larralde, António J. M. Ribeiro, Georg Zeller, and Janet M. Thornton. Investigating enzyme function by geometric matching of catalytic motifs. bioRxiv, 2026.

7. John Jumper, Richard Evans, Alexander Pritzel, Tim Green, Michael Figurnov, Olaf Ronneberger, Kathryn Tunyasuvunakool, Russ Bates, Augustin Žídek, Anna Potapenko, et al. Highly accurate protein structure prediction with alphafold. nature, 596(7873):583–589, 2021.

8. Christopher P. Landowski, Anne Huuskonen, Ramon Wahl, Ann Westerholm-Parvinen, Anne Kanerva, Anna-Liisa Hänninen, Noora Salovuori, Merja Penttilä, Jari Natunen, Christian Ostermeier, Bernhard Helk, Juhani Saarinen, and Markku Saloheimo. Enabling low cost biopharmaceuticals: A systematic approach to delete proteases from a well-known protein production host trichoderma reesei. 10(8):e0134723, 2015. Publisher: Public Library of Science.

9. Yu Li, Qinghao Wei, Mohd Sadeeq, Shibin Cui, Jia Zuo, and Peng Xiong. Unlocking the potential of trichoderma reesei as a super-host for heterologous protein production: Challenges, advances, and perspectives. 20(9):e70121, 2025.

10. Joao PA Moraes, Gisele L Pappa, Douglas EV Pires, and Sandro C Izidoro. Gass-web: a web server for identifying enzyme active sites based on genetic algorithms. Nucleic acids research, 45(W1):W315–W319, 2017.

11. OpenAI. Chatgpt (gpt-5.1), 2026. Large language model.

12. Verena Seidl and Bernhard Seiboth. Trichoderma reesei: genetic approaches to improving strain efficiency. 1(2):343–354, 2010. Publisher: Taylor & Francis eprint: 10.4155/bfs.10.1.

13. Karel van der Weg, Erinc Merdivan, Marie Piraud, and Holger Gohlke. Topec: prediction of enzyme commission classes by 3d graph neural networks and localized 3d protein descriptor. Nature Communications, 16(1):2737, Mar 2025.

14. Yuxin Yang, Abby Jerger, Song Feng, Zixu Wang, Christina Brasfield, Margaret S. Cheung, Jeremy Zucker, and Qiang Guan. Improved enzyme functional annotation prediction using contrastive learning with structural inference. Communications Biology, 7(1):1690, Dec 2024.

15. Tianhao Yu, Haiyang Cui, Jianan Canal Li, Yunan Luo, Guangde Jiang, and Huimin Zhao. Enzyme function prediction using contrastive learning. Science, 379(6639):1358–1363, 2023.

16. Tianhao Yu, Haiyang Cui, Jianan Canal Li, Yunan Luo, Guangde Jiang, and Huimin Zhao. CLEAN: git repository. https://github.com/tttianhao/CLEAN, 2025. Accessed: 2026-02-18.

17. Chengxin Zhang, Lydia Freddolino, and Yang Zhang. Cofactor: improved protein function prediction by combining structure, sequence and protein–protein interaction information. Nucleic acids research, 45(W1):W291–W299, 2017.

18. Yang Zhang and Jeffrey Skolnick. Tm-align: a protein structure alignment algorithm based on the tm-score. Nucleic acids research, 33(7):2302–2309, 2005.

